# Darwin21 Genome Database: A Curated Whole-Genome Repository of Endophytic Bacteria from Desert Plants

**DOI:** 10.1101/2025.08.25.672100

**Authors:** Sabiha Parween, Arun Prasanna Nagarajan, Amal K. Alghamdi, Abdul Aziz Eida, Feras F. Lafi, Luma Albanna, Nida Salem, Barakat Abu-Irmaileh, Zaid A. Pirzada, Shahid Siddique, Ruben Garrido-Oter, Paul Schulze-Lefert, Maged. M. Saad, Heribert Hirt

## Abstract

Microbial communities associated with desert plants play a pivotal role in enhancing host survival under extreme environmental stressors, including drought, salinity, and nutrient limitation. The Darwin21 Endophytic Microbial Collection is one of the largest curated repositories of 2,500 cultivable endophytic bacteria isolated from 23 native desert plant species across Saudi Arabia, Jordan, and Pakistan. Representing a broad spectrum of arid microhabitats from inland deserts and mountain wadis to coastal mangroves and date palm oases, the collection supports integrative studies on microbial ecology and plant–microbe interactions in water-limited ecosystems. A central component of this initiative is the Darwin21 Genome Database, which currently hosts whole-genome sequences (WGS) of 534 endophytic bacterial isolates annotated with extensive ecological metadata, assembly statistics, functional traits, and host associations. The database interface provides tools for genome exploration, metadata filtering, and functional gene mining, enabling users to identify taxa and traits of agronomic interest, particularly for applications in sustainable agriculture and sustainable desert revegetation. By combining genomic, ecological, and functional data, the Darwin21 Genome Database serves as a foundational platform for the development of targeted microbial inoculants and fosters data-driven research into desert microbiomes and plant resilience mechanisms.

**Database URL:** https://www.genomedatabase.org/

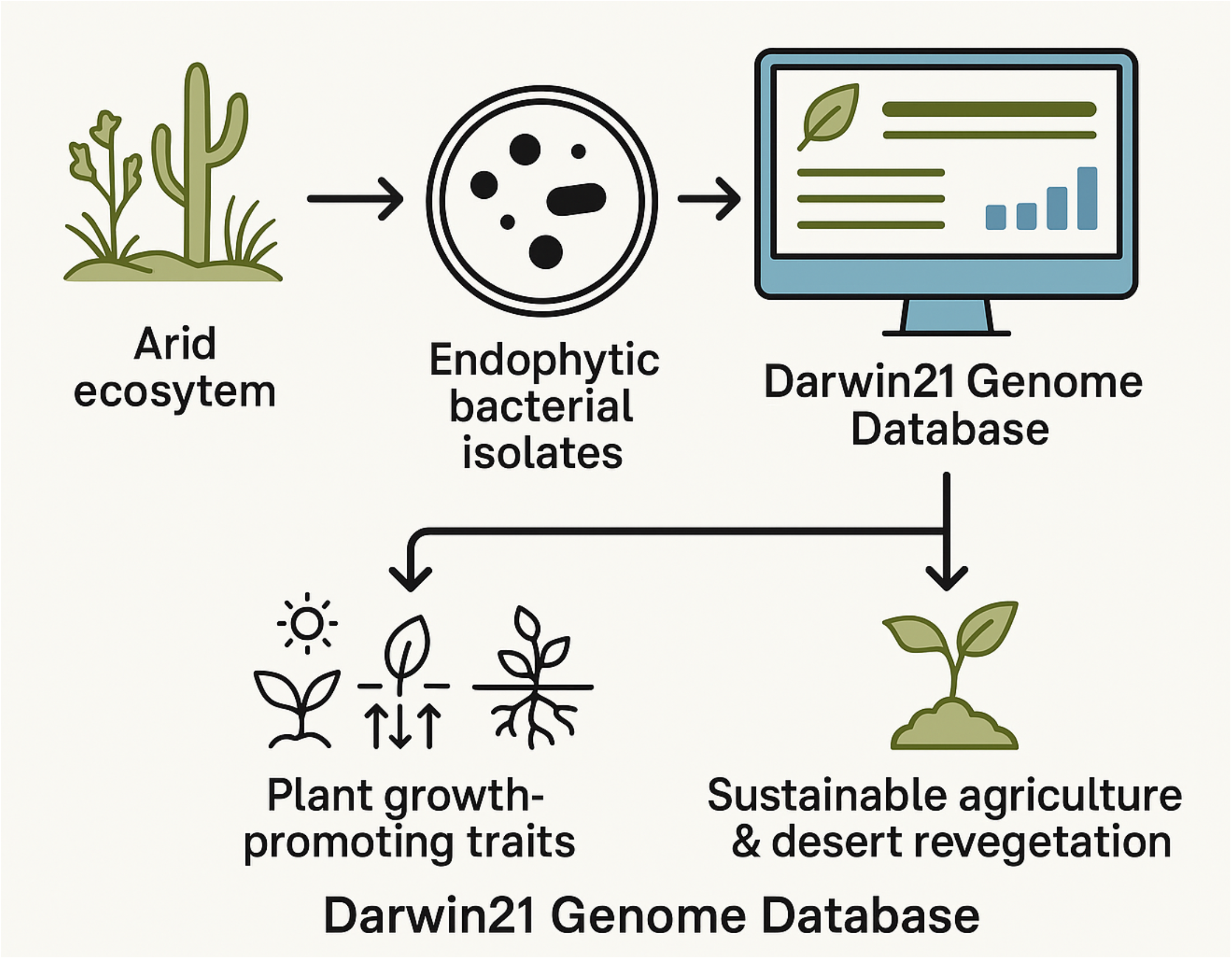

## Introduction

Desert ecosystems represent some of the most challenging environments for plant life, characterized by intense heat, limited water availability, and high salinity. Despite these conditions, many native plant species have evolved physiological and biochemical strategies to persist and thrive. An emerging body of research has highlighted the critical role of plant-associated microbes, particularly endophytic bacteria, in mediating plant adaptation to such extreme abiotic stressors (Alsharif et al., 2020; van der Lelie et al., 2009). These microbes, which reside within plant tissues without causing harm, contribute to host fitness by enhancing stress tolerance, facilitating nutrient acquisition, and synthesizing phytohormones and other growth-promoting compounds (Saad et al., 2020).

The Darwin21 project was launched to systematically investigate and catalogue the endophytic microbial communities associated with native desert plants, with a specific focus on isolating strains with beneficial traits relevant to sustainable agriculture. By conducting extensive field sampling across arid and semi-arid habitats, including inland deserts, saline coastal zones, and mountainous wadis, the project has assembled one of the most comprehensive collections of cultivable endophytic bacteria from desert flora to date (Bang et al., 2018). As a central outcome of this initiative, we introduce the Darwin21 Genome Database, a curated and publicly accessible resource containing whole-genome sequences (WGS) from over 500 bacterial strains. These strains were isolated from 23 ecologically diverse desert plant species, encompassing a broad taxonomic range that includes nitrogen-fixing legumes, halophytes, and pioneer grasses native to the country isolated from. The sequenced isolates span several major bacterial phyla, including Proteobacteria, Actinobacteria, Firmicutes, and Bacteroidetes, reflecting the taxonomic and ecological breadth of the Darwin21 collection.

Each genome entry in the database is enriched with detailed contextual metadata, including plant host identity, geographic origin, isolation conditions, and functional annotations. Of particular interest are predicted plant growth-promoting traits (PGPTs) (Alghamdi et al., 2023), such as genes associated with auxin biosynthesis, phosphate solubilization, and stress response. This enables users to perform genome mining for agricultural applications, comparative genomic analysis, and exploration of microbial adaptations to arid environments (Eida et al., 2020; Andrés-Barrao et al., 2017; Lafi et al., 2017a, Lafi et al., 2017b; (Alghamdi et al., 2024 a, Alghamdi et al., 2024 b)).

By integrating ecological metadata with high-quality genomic and functional annotations, the Darwin21 Genome Database offers a resource platform for research in microbial ecology, host-microbe interactions, and climate-resilient crop development. We describe the structure, content, and navigation features of the database, and discuss its utility for researchers aiming to identify, compare, and deploy beneficial microbes for use in sustainable agriculture.

## Materials and Methods

### Selection of Plant Hosts

The selection of plant hosts for the Darwin21 endophytic bacterial collection was guided by a comprehensive review of ecological literature and targeted field surveys across arid and semi-arid landscapes. A total of 23 native plant species were chosen to represent a wide range of ecological functions and environmental adaptations. These species were selected for their prevalence in desert ecosystems, their resilience to abiotic stresses such as drought and salinity, and their ecological roles in soil stabilization, nutrient cycling, and vegetation recovery. The sampling sites encompassed diverse habitats, including inland deserts, saline coastal plains, and mountainous wadis across Saudi Arabia and adjacent regions.

To ensure broad phylogenetic coverage, the selected plant species were distributed across two major botanical classes, Magnoliopsida (17 species) and Liliopsida (6 species), spanning 10 botanical orders and 12 plant families. Key families included Fabaceae, Poaceae, Amaranthaceae, and Asteraceae, which are known to harbor diverse and functionally relevant microbial partners. The 23 species encompassed 19 genera, providing a diverse genetic and ecological foundation for microbial isolation **(Supplementary Table 1)**.

The ecological characteristics of the host plants were used as a contextual framework for selecting species likely to harbor stress-adapted microbial communities. Leguminous plants such as *Acacia gerrardii, Indigofera argentea*, and *Astragalus tribuloides* were included due to their capacity for symbiotic nitrogen fixation (Graham and Vance, 2003; Vitousek et al., 2002), which is essential in nutrient-depleted soils. Halophytic species like *Avicennia marina* (Alghamdi et al., 2024(b)), *Halothamnus bottae*, and *Zygophyllum simplex* were selected for their natural tolerance to salinity, offering insights into microbial adaptations in hypersaline environments. Pioneer species and soil-stabilizing grasses such as *Panicum turgidum, Dactyloctenium aristatum*, and *Erodium* spp. were chosen for their role in ecological succession and erosion control. In addition, the culturally and economically significant date palm, *Phoenix dactylifera* (Ajwa cultivar), was included for its long-standing association with desert agriculture and its potential as a model system for microbiome research (Shivanandappa et al., 2023; Cherif et al., 2015).

### Microbial Isolation and Identification

Root and shoot tissues were surface-sterilized and processed using standardized microbiological techniques as described on (Alghamdi et al., 2024 a, Alghamdi et al., 2024b); Lafi et al., 2016; Alghamdi et al., 2023). Culturable bacterial endophytes were isolated under sterile conditions, using a serial dilution technique. The serial dilutions were then plated onto different media for optimal bacterial isolation this includes Luria-Bertani (LB), Yeast extract (L7025, Sigma-Aldrich), Tryptic Soy Agar (TSA) and Zobell Marine (ZM) Agar. The purified bacteria were preserved in glycerol stocks for long-term storage. Strains were preliminarily characterized using phenotypic assays and 16S rRNA gene sequencing. A selected subset of strains underwent whole-genome sequencing to facilitate detailed functional and taxonomic analysis (Bang et al., 2018).

### Genomic Sequencing, Assembly, Annotation and Phylogeny

Genomic DNA (gDNA) was extracted from pure bacterial cultures grown on LB agar plates incubated at 30 °C for 48 hours. Total gDNA was isolated using the GenElute*™* Bacterial Genomic DNA Kit (Sigma-Aldrich, Germany) according to the manufacturer’s instructions. DNA quality and concentration were assessed using spectrophotometry and agarose gel electrophoresis. Sequencing libraries were prepared using multiple platforms, including Illumina HiSeq, HiSeq 3000, MiSeq, PacBio RSII, and PacBio Sequel instruments. Illumina sequencing was performed using the Nextera XT DNA Library Prep Kit (or equivalent), producing 150 bp paired-end reads. PacBio libraries were constructed using the SMRTbell Template Prep Kit. Raw Illumina reads underwent quality control using fastp v0.20.1 (Chen et al., 2018), while PacBio reads were processed using Guppy or the SMRT Analysis Suite, with further filtering by Filtlong v0.2.1.

Genome assemblies were generated using different pipelines depending on sequencing technology: A4Pipeline (20160825), SPAdes (v3.6–v3.15.5) (Bankevich et al., 2012) for Illumina reads, and HGAP (v4), smrtpipe v2.3.0.140936.p5, or HGAP.2 (v2.3.0) for PacBio data. Assembly quality metrics, including the number of contigs, N50 values, genome size, GC content, and completeness, were computed using QUAST v5.0.2 (Gurevich et al., 2013) and CheckM v1.2.2 (Parks et al., 2015). The average genome completeness exceeded 95%, with contamination generally below 3%.

Genome assemblies annotated using Prokka with predicted proteins (^*^.faa) were analyzed with Roary v3.xx (Page et al., 2015) to identify core genes. Core gene alignments were generated using MAFFT v7.xx (Katoh & Standley, 2013), trimmed with trimAl v1.4 (Capella-Gutiérrez et al., 2009), and used to construct a maximum-likelihood phylogenetic tree with FastTree v2.xx (Price et al., 2010) under the GTR+Gamma model. The tree was visualized in iTOL (Letunic & Bork, 2021).

Functional annotation of assembled genomes was performed using Prokka v1.14.6 (Seemann, 2014). KEGG Orthology (KO) terms were assigned using eggNOG-mapper v2.1 (Huerta-Cepas et al., 2017), facilitating downstream functional analysis.

### Gene mining for potential plant growth-promoting traits

To functionally characterize plant-beneficial traits, we utilized the Plant-associated Bacteria (PLaBAse) resource (Patz et al., 2021), which curates over 6.9 million proteins representing 6,900 unique PGPTs. Protein-level annotations were assigned using a combined BLASTp + HMMER approach against the PGPT database. Hits were filtered with a threshold of ≥40% identity and ≥70% query coverage to ensure confident trait assignments. This allowed systematic profiling of PGPTs across strains into Direct Effects, Indirect Effects, and Putative PGPTs.

### Database construction

To effectively store, manage, and deliver genomic and functional data, a MySQL relational database management system was employed. Custom database schema designs were implemented to ensure referential integrity across multiple tables encompassing genomic sequences, functional annotations, ecological metadata, and plant growth-promoting trait (PGPT) classifications. Schema normalization and indexing strategies were used to optimize query performance and facilitate scalable data retrieval (Elmasri & Navathe, 2015). A custom WordPress-based front-end was developed to allow seamless integration between the database and user-facing components. PHP scripts and templating logic dynamically rendered genome-specific records and facilitated secure downloads of associated files, including FASTA, GFF, and protein annotations. The interface leveraged the Data Tables JavaScript library for building rich, client-side interactivity, including auto-complete search, column-specific filtering, pagination, and dynamic table rendering for fast browsing of large datasets (Allan et al., 2022). The graphical user interface (GUI) was built using standard web technologies: HTML5 for semantic content structure, CSS3 for responsive styling, and JavaScript for asynchronous interactions. These tools enabled a clean, responsive design that functions effectively across modern browsers and devices. For hosting, the platform is deployed on a Virtual Private Server (VPS) with secure HTTPS access, ensuring high availability and rapid response times under concurrent user loads (**Supplementary Figure 1**). Server-side caching and load-balancing mechanisms were employed to support smooth data access, even with increasing traffic volumes.

## Results and Discussion

### Database structure and navigation

The Darwin21 Genome Database is organized as a genome-centric platform featuring multiple interactive sections and tools for exploring microbial data associated with desert plant endophytes, as shown in **Figure 1**. The resource integrates comprehensive genome records that include detailed assembly metrics such as scaffold counts, N50 values, genome size, and assembly methods, alongside quality assessments based on CheckM lineage assignments and completeness estimates (Parks et al., 2015). Each entry links these data with extensive ecological metadata, including host plant information, country of isolation, sequencing platform, and accession numbers. Users can access and download a range of annotation files, including GFF annotations, predicted protein sequences, and nucleotide sequences.

**Figure 1.**
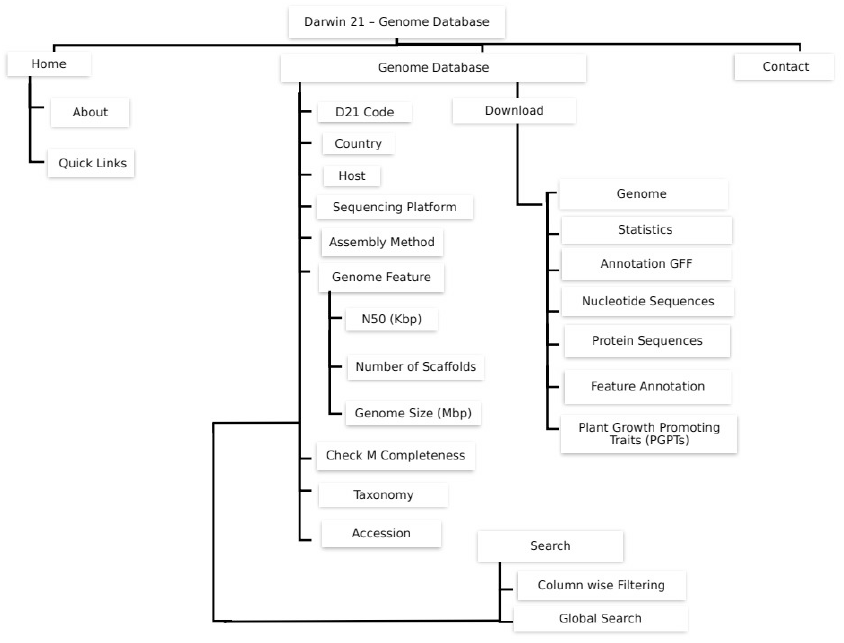
Overview of the Darwin21 Genome Database architecture. The database is organized around genome-centric records, integrating assembly statistics, ecological metadata, functional annotations, and plant growth-promoting trait (PGPT) profiles. The main navigation includes a home page with summary views, a quick links panel for accessing downloads, statistics, and search functions, and detailed genome pages showing assembly information (scaffold count, genome size, N50, etc.), CheckM-based quality metrics, and source metadata such as host plant, country, and sequencing platform. Users can interactively filter and sort data, perform global searches, and access downloadable annotation files (GFF, protein, nucleotide, etc). The interface supports dynamic exploration of taxonomy, gene content, and functional traits across the database.

The database provides integrated feature annotations covering taxonomy, genome features, and plant growth-promoting traits (PGPTs) (Patz et al., 2021). The interface emphasizes interactive tables where users can sort columns by attribute, filter records by values such as host plant, country, or sequencing platform, and download genome data and associated annotations directly. Annotation and metadata are linked, enabling seamless navigation from genome assembly details to functional profiles and PGPT information. Contact and help options are always accessible from the main navigation panel. An intuitive navigation system supports column-wise filtering and global keyword searches across all records, enabling efficient retrieval and comparison of genome entries. Interactive tables allow users to sort and filter data dynamically and to move seamlessly between genome assembly details and functional annotations. Additionally, **Supplementary Table 1** presents the taxonomic classification and ecological role of each plant host included in the study. Also the whole-genome phylogenetic relationships among strains were inferred at the protein level from concatenated core gene alignments (Supplementary Figure 2), providing high-resolution insights into strain relatedness, clonal clustering, and overall genomic diversity.

### Genome-Based Access to Potential Plant Beneficial Traits

We systematically categorized microbial genes into distinct functional classes relevant to plant growth and stress resilience. Each microbial strain was profiled based on its encoded PGPTs and visualized through pie charts summarizing three major trait categories: Direct Effects, Indirect Effects, and Putative PGPTs and its related subcategories

Direct Effects encompass traits that directly enhance plant performance. These include genes involved in bio-fertilization processes such as nitrogen fixation, phosphate and potassium solubilization, sulfur assimilation (Vessey, 2003; Richardson et al., 2009), as well as carbon dioxide fixation (Lugtenberg & Kamilova, 2009). Additionally, pathways contributing to phytohormone biosynthesis or modulation—including auxins, gibberellins, cytokinins, and ethylene (Spaepen et al., 2007; Dodd et al., 2010) were identified. Additionally, microbial pathways involved in promoting plant development through signals such as root and shoot growth stimulation, branching, germination, and vitamin production were included. Indirect Effects include gene functions that enhance plant resilience through the mitigation of biotic and abiotic stressors. These involve biosynthesis of antimicrobial compounds, detoxification of heavy metals and xenobiotics, and activation of plant immune responses, including induced systemic resistance (ISR) and systemic acquired resistance (SAR) van Loon et al., 1998; Pieterse et al., 2014). The dataset also includes traits that facilitate microbial colonization of plant tissues, including chemotaxis, motility, surface attachment, and enzymatic degradation of plant-derived substrates. (Compant et al., 2010; Hardoim et al., 2015). Putative PGPTs represent genes with the potential to play plant-beneficial roles that currently lack experimental validation. These were retained to ensure a broad functional inventory that supports hypothesis generation and further exploration. All annotated functional profiles, along with interactive pie chart visualizations for each strain, are accessible for browsing and download through the Darwin21 database.

### Darwin 21-a valuable and pioneering genomic resource

Darwin21 represents a unique and forward-looking genomic repository that integrates ecological context with high-resolution functional genomic data, with specific emphasis on microbial endophytes associated with desert plants. Unlike conventional databases, Darwin21 interlinks high-quality genome assemblies with rich ecological metadata and experimentally relevant annotations, offering a comprehensive platform for investigating plant-microbe interactions in arid and extreme environments. Its trait-centric framework enables the identification of plant growth-promoting genes across both experimentally validated and putative functional categories, providing unprecedented granularity for functional and ecological interpretation. The platform’s integration of interactive tools, customizable search queries, and user-friendly visualizations empowers researchers to derive biologically meaningful insights efficiently. As global interest in sustainable agriculture intensifies and climate-resilient crop systems continue to grow, Darwin21 delivers critical infrastructure to support the exploration and application of microbial solutions aimed at enhancing plant resilience, nutrient acquisition, and environmental adaptation under abiotic stress conditions.

## Conclusions

In conclusion, Darwin21 emerges as a critical platform for unlocking the genomic and functional potential of desert plant-associated microbiomes. By integrating high-quality genome data with ecologically meaningful annotations, it enables the systematic identification of plant-beneficial traits across diverse microbial taxa. Its interactive tools and trait-based framework not only facilitate in-depth exploration but also bridge the gap between basic microbial genomics and applied agricultural innovation. As the demand for climate-resilient and sustainable farming solutions grows, Darwin21 will support future research and practical applications that harness the power of microbiomes to enhance crop performance under environmental stress.

## Author contributions

HH and MMS designed the project. HH, MMS, AKA, LA, NS, FLL, ZAP, AAE and SS collected the material and cultivated the strains,. BAI determined the plant species. After sequencing, SP, AN, and RGO did the bioinformatic analysis. SP develops the design and structure of the database. SP, MMS, and HH contributed to the paper writing.

## Acknowledgment

We gratefully acknowledge the Darwin21 team for their valuable technical assistance and insightful discussions throughout this study. We thank Olga Artyukh, Hanna Cherviakouskaya, Busra Elkatmis, Yao Wu, Wiam Alshari, Rewaa Jalal, Linah Alghanmi, Khairiah Alwutayd, Fatimah Abdulhakim, Eneroliza Vande Loo and Carlos A. Penaloza Rodriguez for their dedicated support, which greatly contributed to the quality and consistency of the work.

## Data availability

Most of the strains’ genomic data have also been submitted to NCBI, under the projects PRJNA1042672, PRJNA1054060, PRJNA1076458, PRJNA312745, PRJNA313373, PRJNA313377, PRJNA345401, PRJNA352128, PRJNA398615, PRJNA708169, PRJNA740144, PRJNA827473, PRJNA929179, PRJNA973967. The data underlying this article can be accessed at **https://www.genomedatabase.org/**

## Funding

The work was funded by the KAUST fund BAS/1/1062-01-01 to HH as part of the DARWIN21 desert initiative (http://www.darwin21.org/).

## Conflict of interest

The authors declare that the research was conducted in the absence of any commercial or financial relationships that could be construed as a potential conflict of interest.

## Consent for Publication

There is no conflict of consent for publication

## Code availability

The code avaiability for database construction is available at Supplementart File 1.

---

**Supplementary Figure S1.**
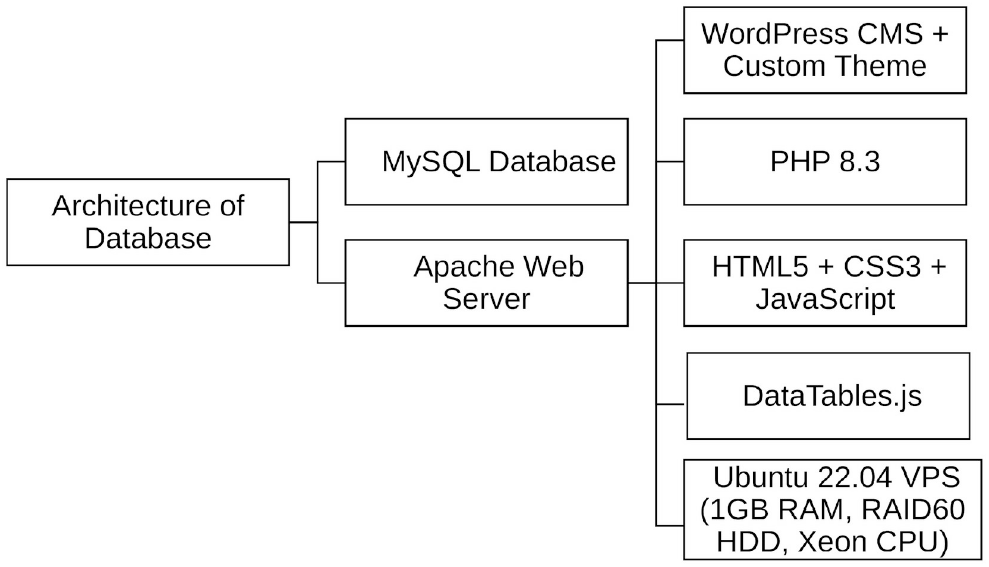
System architecture of the Darwin21 Genome Database. The database infrastructure comprises a MySQL relational database for storing genomic, annotation, and metadata records; an Apache web server integrated with WordPress CMS using a custom theme and PHP 8.3 for dynamic content generation. The front end is designed using HTML5, CSS3, and JavaScript, with interactive tables powered by the DataTables.js library. The platform is deployed on a Virtual Private Server (VPS) running Ubuntu 22.04 with 1GB RAM, RAID60 HDD, and Xeon CPU configuration, ensuring scalable performance and high availability.

**Supplementary Figure 2.**
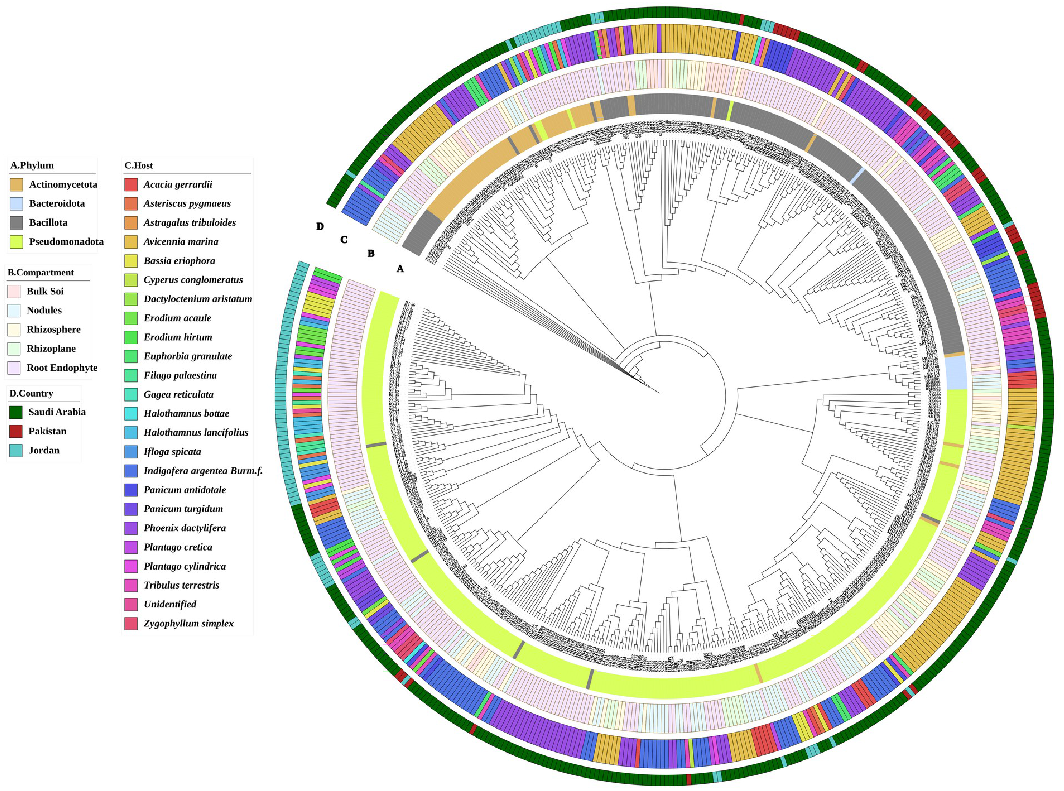
Maximum-likelihood phylogenetic tree of microbial strains. The tree is annotated with four categorical layers: **A** (Phylum), **B** (Compartment), **C** (Plant host), and **D** (Country). Category A is color-coded according to bacterial phyla identified by taxon profile of assembled sequences. Category B distinguishes the sample compartments. Category C denotes the plant host species from which each sample was obtained, and category D represents the country of sample origin. Branch lengths indicate evolutionary distances, and bootstrap support values (>90%) are not displayed to maintain clarity. The arrangement of colored symbols adjacent to each leaf allows rapid visual assessment of taxonomic affiliation, ecological context, and geographic provenance within the phylogenetic framework.

**Supplementary Table 1:** Taxonomy and Ecological traits of each hos.

| Scientific_Name | Genus | Family | Order | Class | Phylum | Kingdom | Ecological_Role |
| --- | --- | --- | --- | --- | --- | --- | --- |
| *Acacia gerrardii* | Acacia | Fabaceae | Fabales | Magnoliopsida | Tracheophyta | Plantae | Nitrogen fixer, soil stabilizer, fodder source |
| *Asteriscus pygmaeus* | Asteriscus | Asteraceae | Asterales | Magnoliopsida | Tracheophyta | Plantae | Annual herb, pioneer species in disturbed desert habitats |
| *Astragalus tribuloides* | Astragalus | Fabaceae | Fabales | Magnoliopsida | Tracheophyta | Plantae | Nitrogen-fixing legume in arid grasslands |
| *Avicennia marina* | Avicennia | Acanthaceae | Lamiales | Magnoliopsida | Tracheophyta | Plantae | Mangrove species, shoreline stabilizer, carbon sequestration |
| *Bassia eriophora* | Bassia | Amaranthaceae | Caryophyllales | Magnoliopsida | Tracheophyta | Plantae | Salt-tolerant shrub, indicator of saline soils |
| *Cyperus conglomeratus* | Cyperus | Cyperaceae | Poales | Liliopsida | Tracheophyta | Plantae | Sand binder in arid ecosystems |
| *Dactyloctenium aristatum* | Dactyloctenium | Poaceae | Poales | Liliopsida | Tracheophyta | Plantae | Grass species aiding in dune stabilization |
| *Erodium acaule* | Erodium | Geraniaceae | Geraniales | Magnoliopsida | Tracheophyta | Plantae | Annual herb, contributes to early soil coverage |
| *Erodium hirtum* | Erodium | Geraniaceae | Geraniales | Magnoliopsida | Tracheophyta | Plantae | Annual herb, pioneer species in degraded lands |
| *Euphorbia granulate* | Euphorbia | Euphorbiaceae | Malpighiales | Magnoliopsida | Tracheophyta | Plantae | Drought-adapted herb, potential allelopathic effects |
| *Filago palaestina* | Filago | Asteraceae | Asterales | Magnoliopsida | Tracheophyta | Plantae | Ephemeral plant, desert soil cover |
| *Gagea reticulata* | Gagea | Liliaceae | Liliales | Liliopsida | Tracheophyta | Plantae | Bulbous geophyte, early-season desert flora |
| *Halothamnus bottae* | Halothamnus | Amaranthaceae | Caryophyllales | Magnoliopsida | Tracheophyta | Plantae | Halophyte, adapted to extreme arid saline soils |
| *Halothamnus lancifolius* | Halothamnus | Amaranthaceae | Caryophyllales | Magnoliopsida | Tracheophyta | Plantae | Halophyte, common in degraded saline areas |
| *Ifloga spicata* | Ifloga | Asteraceae | Asterales | Magnoliopsida | Tracheophyta | Plantae | Small annual herb, colonizer in sandy soils |
| *Indigofera argentea* Burm.f. | Indigofera | Fabaceae | Fabales | Magnoliopsida | Tracheophyta | Plantae | Nitrogen-fixing legume, improves poor soils |
| *Panicum antidotale* | Panicum | Poaceae | Poales | Liliopsida | Tracheophyta | Plantae | Perennial grass, erosion control, forage species |
| *Panicum turgidum* | Panicum | Poaceae | Poales | Liliopsida | Tracheophyta | Plantae | Dominant perennial grass in |

## Notes

https://www.genomedatabase.org/

### Competing Interest Statement

The authors have declared no competing interest.

