## Supplemental Table 1 for "Darwin21 Genome Database: A Curated Whole-Genome Repository of Endophytic Bacteria from Desert Plants"

### Code Availability and Implementation Details: Genome Database Table (WordPress + DataTables)

**Date:** August 11, 2025

#### Overview

This document describes the availability, environment, and implementation details for the **Genome Database** table view built in WordPress using PHP (wpdb), Bootstrap, and jQuery DataTables. It is intended for inclusion in the manuscript's **Code Availability** section and for reproducibility by other researchers.

#### Code Location and Access

- **Framework:** WordPress (PHP)

- **Primary components:** Theme/Plugin PHP template rendering a Bootstrap table, client-side filtering and rendering with jQuery DataTables.

- **Table source:** WordPress MySQL table `{$wpdb->prefix}genomes`.

- **Static file base path:** `https://www.genomedatabase.org/Genome/536_ASSEMBLED_GENOMES/` (used to build links for `.fna`, `.gff`, `.faa`, `.ffn`, `.txt`, `.tsv`).

If a repository URL is available, include it here. Otherwise, the full code snippet is embedded below (Appendix A).

#### System Requirements

- **Server:** Linux (tested on Ubuntu 20.04+), Apache/Nginx with PHP 7.4+ (PHP 8.x compatible)

- **CMS:** WordPress 6.x

- **Database:** MySQL/MariaDB (as used by WordPress)

- **Front-end libraries:**

- Bootstrap CSS: `admin/css/bootstrap.min.css` within the active theme

- jQuery (bundled by WordPress)

- DataTables (JS/CSS; enqueued or loaded via theme/plugin)

#### Installation & Deployment

1. Create database table `wp_genomes` (or `{prefix}genomes`) with columns used by the snippet (e.g., `ID, D21_Genome, No_Scaffolds, N50_Kbp, GenomeSize_Mbp, CheckM_Lineage, CheckM_Completeness, Host, Isolation_source, Country, Assembly, Sequencing_Technologies, Kingdom, Phylum, Class, Order, Family, Genus, Bioproject, Accessions_Biosample`).

2. Place the PHP/HTML/JS snippet into a theme template or a custom plugin page hooked into WordPress admin or a front-end page.

3. Ensure Bootstrap and DataTables are enqueued (or include the CSS/JS links) before this template runs.

4. Set the base path for assembled genomes:

- `$base_path = 'https://www.genomedatabase.org/Genome/536_ASSEMBLED_GENOMES/';`

5. Verify file naming pattern on the server:

- For each row where `$folder = $genome->D21_Genome`, the code links:

- `{base}/{folder}/{folder}.fna` (Genome)

- `{base}/{folder}/{folder}.gff` (GFF)

- `{base}/{folder}/{folder}.faa` (Protein)

- `{base}/{folder}/{folder}.ffn` (Nucleotide)

- `{base}/{folder}/{folder}.txt` (Statistic)

- `{base}/{folder}/{folder}.tsv` (Annotation Feature)

#### Usage

- The template queries all genomes:

$genomes = $wpdb->get_results("SELECT * FROM {$wpdb->prefix}genomes", OBJECT);

- Each genome record is rendered as a row with metadata and download links.

- The DataTables header row is cloned to create a **filters** row with **column-specific** text inputs for columns 2–19 (0-based), enabling per-column searching.

- DataTables initialization:

- Horizontal scroll (`scrollX: true`),

- Sticky header (`fixedHeader: true`),

- 15 rows per page (`pageLength: 15`),

- `orderCellsTop: true` for proper sort behavior with dual header rows.

#### Security and Data Integrity

- Sanitize/escape outputs:

- Use `esc_html()` for plain text fields,

- Use `esc_url()` for constructed links,

- Validate `$folder` against an allowlist pattern if needed to prevent path traversal.

- Use **prepared queries** if adding any parameters to the SQL; although current code uses a static `SELECT *`, future filters should use `$wpdb->prepare()`.

- Consider paging on the server side if the table grows large (DataTables server-side mode).

#### Reproducibility Checklist

- WordPress environment and versions (core/plugins/themes) documented

- Database schema exported (SQL) and included

- Sample dataset (e.g., 10 rows) for quick verification

- Static file directory structure mirrors `{base}/{folder}/{folder}.*` pattern

- DataTables and Bootstrap enqueued correctly

- Browser console has no JS errors on load

#### Known Limitations

- Client-side filtering may become slow for very large datasets; consider DataTables server-side processing for >10–20k rows.

- Exact filename dependence: the link builder assumes files exist and match the `{folder}.{ext}` pattern.

- No authentication on downloads: files are public if accessible directly via the constructed URLs (ensure this matches your data policy).

#### How to Cite / Availability Statement (Template)

The WordPress template and JavaScript used to render and filter the Genome Database table are available with this manuscript as supplementary code (Appendix A). The dataset is stored in the WordPress database table `{prefix}genomes`. Pre-assembled genome files referenced by the table are served under the static path `https://www.genomedatabase.org/Genome/536_ASSEMBLED_GENOMES/` and are named as `<D21_Genome>.<ext>` (ext ∈ {{fna, gff, faa, ffn, txt, tsv}}). The code is released under an open-source license; please include license terms if required by your journal.

#### Appendix A — Full Code Snippet

<link rel="stylesheet" href="<?php echo get_stylesheet_directory_uri(); ?>/admin/css/bootstrap.min.css">

<style>.genomecard {{ max-width:100%; }}</style>

<div class="genomeswrap">

<div class="container-fluid">

<div class="row mb-4">

<div class="col-12">

<div class="card genomecard p-2">

<div class="card-body">

<h4>Genome Database</h4>

<table id="myTable" class="table table-striped table-bordered w-100">

<thead>...</thead>

<tbody>

<?php

global $wpdb;

$genomes = $wpdb->get_results("SELECT * FROM {$wpdb->prefix}genomes", OBJECT);

$base_path = 'https://www.genomedatabase.org/Genome/536_ASSEMBLED_GENOMES/';

foreach ($genomes as $genome) { $folder = $genome->D21_Genome; ?>

<tr>

<td><?php echo $genome->ID; ?></td>

<!-- more cells -->

<td><a href="<?php echo $base_path . $folder . '/' . $folder . '.fna'; ?>" target="_blank">Genome</a></td>

<td><a href="<?php echo $base_path . $folder . '/' . $folder . '.gff'; ?>" target="_blank">GFF</a></td>

<td><a href="<?php echo $base_path . $folder . '/' . $folder . '.faa'; ?>" target="_blank">Protein</a></td>

<td><a href="<?php echo $base_path . $folder . '/' . $folder . '.ffn'; ?>" target="_blank">Nucleotide</a></td>

<td><a href="<?php echo $base_path . $folder . '/' . $folder . '.txt'; ?>" target="_blank">Statistic</a></td>

<td><a href="<?php echo $base_path . $folder . '/' . $folder . '.tsv'; ?>" target="_blank">Annotation Feature</a></td>

</tr>

<?php } ?>

</tbody>

</table>

</div>

</div>

</div>

</div>

</div>

</div>

<script type="text/javascript">

jQuery(document).ready(function($) {

$('#myTable thead tr').clone(false).addClass('filters').appendTo('#myTable thead');

$('#myTable thead tr.filters th').each(function (i) {

if (i >= 20 || i <= 1) { $(this).html(''); }

else {

$(this).html('<input type="text" placeholder="Search" style="width:100%; box-sizing:border-box;" />');

$('input', this).on('keyup change', function () {

if ($('#myTable').DataTable().column(i).search() !== this.value) {

$('#myTable').DataTable().column(i).search(this.value).draw();

}

});

}

});

$('#myTable').DataTable({ scrollX:true, fixedHeader:true, pageLength:15, orderCellsTop:true });

});

</script>

*Generated on August 11, 2025.*
